# Class A Scavenger Receptor MARCO negatively regulates Ace expression and aldosterone production

**DOI:** 10.1101/2023.09.26.559578

**Authors:** Conan J O O’Brien, Giorgio Ratti, Hellen Veida-Silva, Emma Haberman, Charles Sweeney, Siamon Gordon, Ana I Domingos

## Abstract

Aldosterone is a potent cholesterol-derived steroid hormone that plays a major role in controlling blood pressure via regulation of blood volume. The release of aldosterone is typically controlled by the renin-angiotensin aldosterone system, situated in the adrenal glands, kidneys, and lungs. Here, we reveal that the class A scavenger receptor MARCO, expressed on alveolar macrophages, negatively regulates aldosterone production and suppresses angiotensin converting enzyme (Ace) expression in the lungs of male mice. Collectively, our findings point to alveolar macrophages as additional players in the renin-angiotensin-aldosterone system and introduce a novel example of interplay between the immune and endocrine systems.

## Introduction

The scavenger receptor MARCO (macrophage receptor with collagenous structure), a class A scavenger receptor, is primarily expressed on macrophages including tumour-associated macrophages, and lung macrophages^1,2^. MARCO, upregulated in response to pathogenic challenge^2^, has been implicated in defence against pathogens in the lung^3^, phagocytosis and clearance of tumor cells ^4^, internalisation of exosomes ^5^, and has been touted as a potential target in anti-cancer immunotherapy ^6^. The class A scavenger receptors have a broad ligand specificity and similar domain structures, comprising a cytoplasmic tail, transmembrane region, spacer region, α helical coiled domain, collagenous domain, and a C-terminal cysteine-rich domain ^7^. Known endogenous ligands for the receptor include, like other class A receptors, modified forms of low-density lipoprotein^7^ indicating that this receptor could be involved in the modulation of cholesterol availability. Scavenger receptors have previously been implicated in corticosteroid output from the adrenal gland. Specifically, Scavenger Receptor BI (SR-BI) was shown to regulate glucocorticoid responses via binding of serum cholesterol, the necessary substrate for adrenal corticosteroids ^8–11^. Aldosterone, the other major adrenal cortex-derived corticosteroid, is known to be regulated by the renin-angiotensin-aldosterone system, which is situated in the adrenals, lung, and kidney.^12^. While a role for scavenger receptors in regulating glucocorticoid responses has been demonstrated, the role of macrophage-expressed scavenger receptors in the regulation of adrenal corticosteroid output at steady state has not yet been explored.

## Results

It is known that statins, cholesterol-lowering drugs have been shown to reduce aldosterone levels in humans ^13,14^. Moreover, *in vitro* studies have demonstrated that cholesterol supplementation boosts the production of aldosterone from cultured cells ^15–17^. Given that the adrenal-derived mouse corticosteroids, most notably corticosterone and aldosterone, derive from cholesterol as the common precursor (Fig. 1a) we hypothesised that cholesterol binding scavenger receptors could modulate adrenal corticosteroid output by regulating the availability of cholesterol that could feed into the steroid hormone biosynthetic pathway. To test this hypothesis, we measured the concentrations of aldosterone and corticosterone in the plasma of Marco^-/-^ and wild-type mice. We found that male Marco^-/-^ mice had significantly elevated levels of plasma aldosterone relative to wild-type (WT) mice (Fig. 1b). In contrast, plasma corticosterone levels were not significantly altered in male mice lacking Marco (Fig. 1c). Marco-deficient female mice did not have altered levels of aldosterone relative to WT, but plasma corticosterone was increased relative to WT counterparts (Fig 1d, e). We observed that the adrenal glands from Marco-deficient male mice were significantly lighter than WT controls in male but not female mice (Fig. 1f, g). To establish whether cholesterol could explain the elevated plasma aldosterone we observe in Marco-deficient male mice, we measured the levels of total serum cholesterol and intra-adrenal cholesterol in males. We found that Marco^-/-^ mice had reduced serum cholesterol relative to WT controls (Fig. 1h), while the normalised levels of intra-adrenal cholesterol were similar between both mouse strains (Fig. 1i). Taken collectively, these findings suggest that, while Marco-deficient male mice have elevated plasma aldosterone concentrations, this is not dependent on systemic or intra-adrenal cholesterol availability. For the purposes of this paper we focused on the phenotype evident in male mice, namely the increase in plasma aldosterone concentrations.

**Figure 1.**
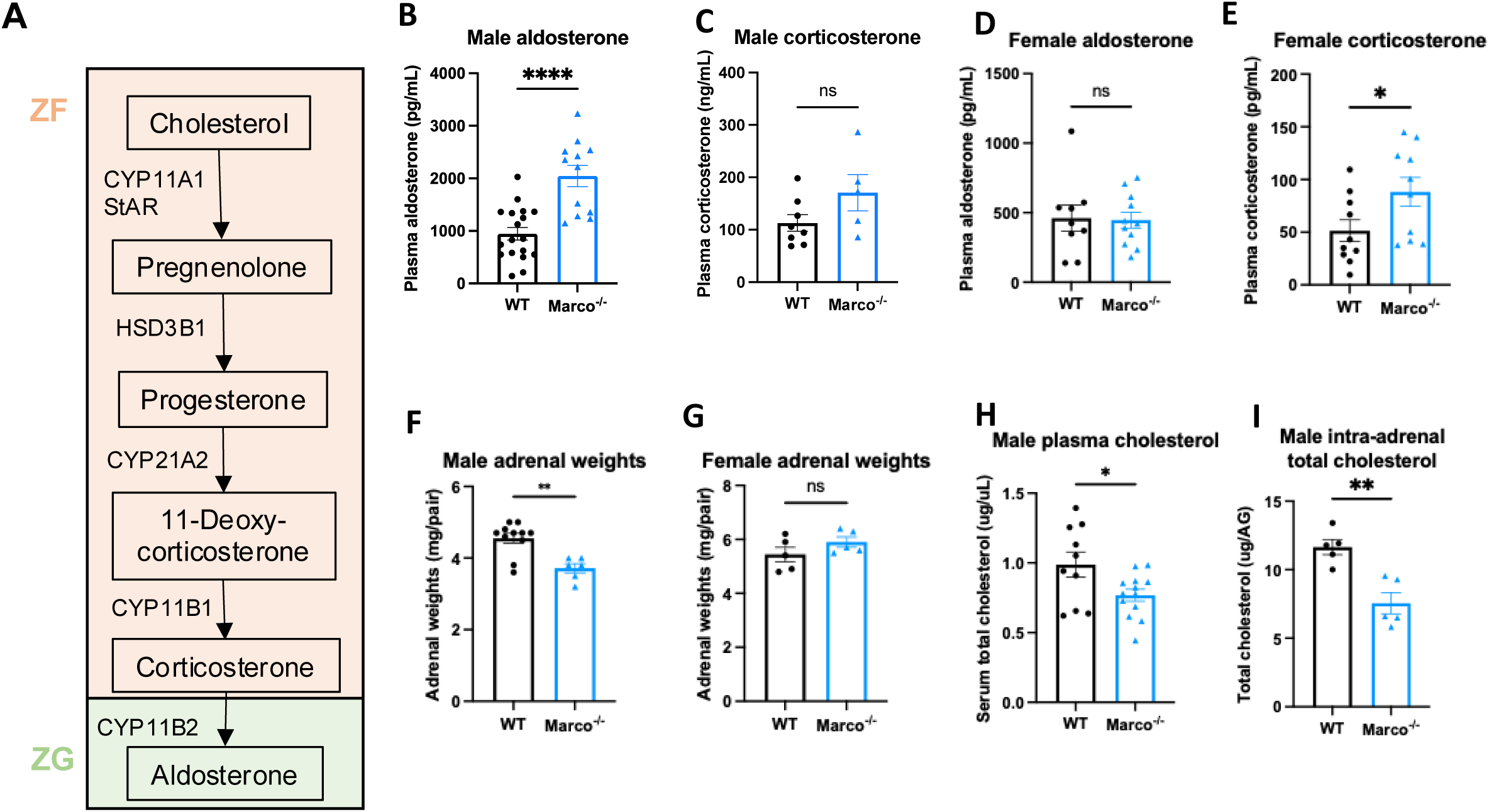
Marco-deficient mice have elevated aldosterone and reduced serum cholesterol. (A) Schematic depicting the murine adrenal corticosteroid biosynthetic pathway. Plasma aldosterone and corticosterone concentrations from wild type (WT) and Marco-deficient male (B, C) and female (D, E) mice as measured by ELISA. (F, G) Weights of both adrenal glands from WT or Marco-deficient male and female mice. Plasma total cholesterol levels (H) and relative intra-adrenal cholesterol levels, normalised to adrenal weight (I), in WT and Marco-deficient male mice. Data in B-H were analysed by two-tailed unpaired Student’s *t*-test and are shown as average ± s.e.m. \**P* < 0.05, \*\**P* < 0.01, \*\*\**P* < 0.001, *\*\*\*\* P* < 0.0001.

We next hypothesised that adrenal gland-derived *Marco* could play a role in modulating aldosterone output. Analysis of publicly available Single cell sequencing data from murine adrenal glands shows that adrenals do contain a substantial population of macrophages expressing *Ptprc* (CD45), *Adgre1* (F4/80), and *Cd68* (CD68). However, we did not detect Marco expression in this cluster of cells, nor any other cluster identified in our analyses (Fig. 2A). This finding was corroborated by immunostaining of male murine adrenal glands, which showed CD68^+^ macrophages in the adrenal zona fasciculata and zona glomerulosa that did not stain positively for MARCO (Fig. 2b). Given that the lung is another site in the RAAS axis, we postulated that Marco-expressing cells in the lung could be involved in mediating the aldosterone phenotype we observed in Fig. 1b. Indeed, single cell RNA seq analysis of the murine lung shows that *Ptprc* (CD45), *Adgre1* (F4/80), and *Cd68* (CD68) expressing cells (Alveolar Macrophages) also express *Marco (*Fig 2c), a finding corroborated by immunostaining in the lung (Fig. 2c). To further validate our single cell sequencing and immunofluorescence data, we carried out qPCR for *Marco* in the lungs and adrenal glands from male WT and Marco^-/-^ mice, which further demonstrated that the lung is the primary site of Marco expression in the RAAS (Fig. 2e).

**Figure 2.**
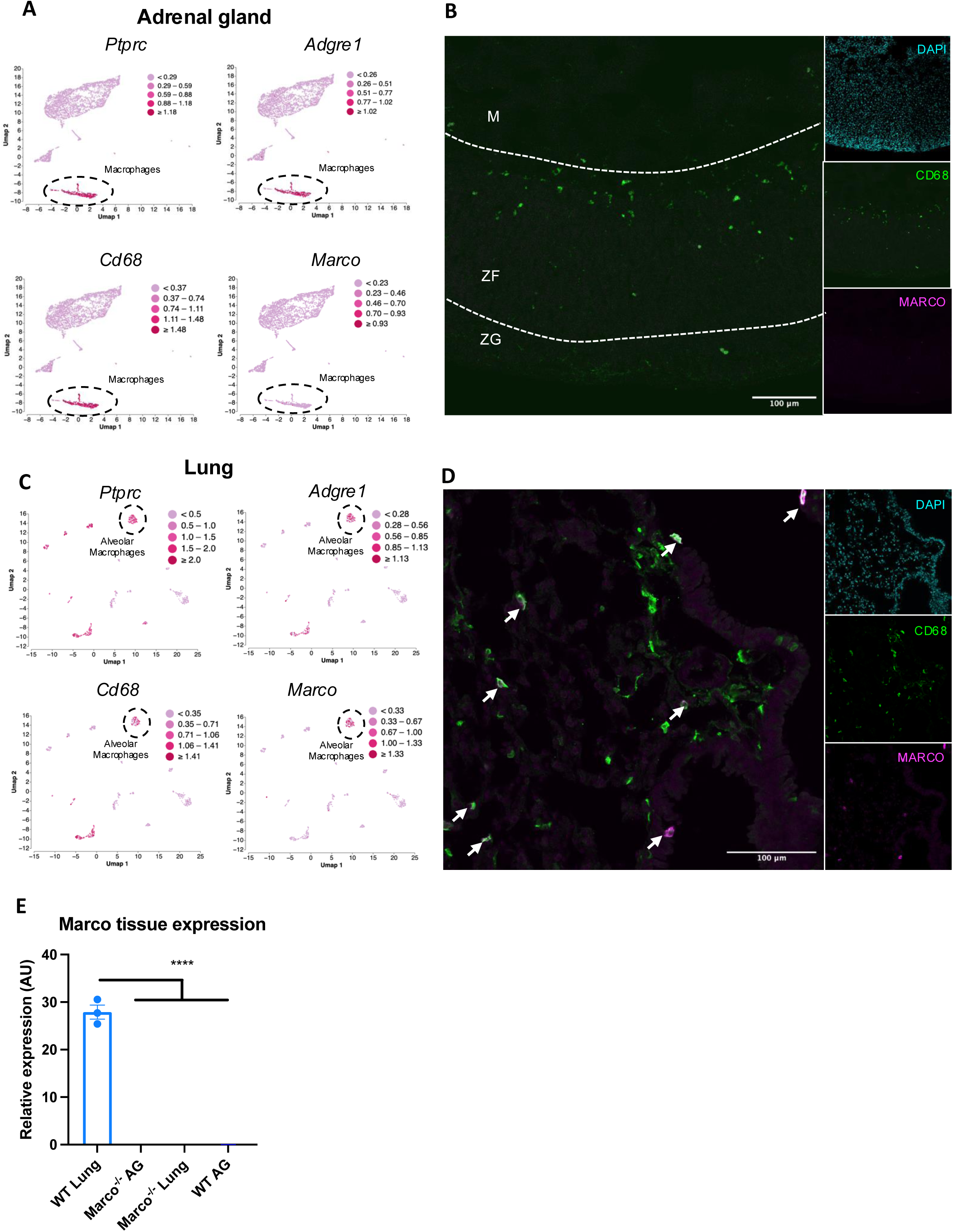
Marco is expressed in the lung and not adrenal glands. (A) Single cell RNA seq data plots from PMID 33571131 representing mRNA expression of Ptprc (CD45), Adgre1 (F4/80), Cd68, and Marco, in male murine adrenal glands. (B) Representative image showing a central cryosection of the male murine adrenal gland from a C57bl/6 mouse stained against CD68 (green) Marco (magenta), and DAPI (cyan). (C) Single cell RNA seq data plots from PMID 30283141 representing mRNA expression of Ptprc (CD45), Adgre1 (F4/80), Cd68, and Marco, in the male murine lung. (D) Representative image showing a cryosection of the male murine lung from a hCD68-GFP reporter mouse stained against GFP (green) Marco (magenta), and DAPI (cyan). (E) qPCR data showing the relative gene expression data for *Marco* in the indicated tissues. M = medulla, ZF = zona fasciculata, ZG = zona glomerulosa.

Aldosterone is a potent blood pressure-regulating hormone, the dysregulation of which can cause severe hypertension and increased cardiovascular risk. It therefore follows that its production is tightly regulated. Aldosterone biosynthesis is fundamentally regulated intra- adrenally by cytochrome P450 family members in the corticosteroid biosynthetic pathway (Fig. 1a). We therefore tested whether altered expression of enzymes in this pathway could explain the hyperaldosteronism observed in Marco-deficient mice. Marco^-/-^ male and female mice showed similar expression of aldosterone biosynthetic enzymes (*Star, Cyp11a1. Hsd3b1, Cyp11b1, Cyp11b2)* as wild type mice (Fig. 3a, b). While *Cyp11b2* (aldosterone synthase) is only expressed in the adrenal zona glomerulosa, other biosynthetic enzymes essential for aldosterone production are expressed in the zona fasciculata.

**Figure 3.**
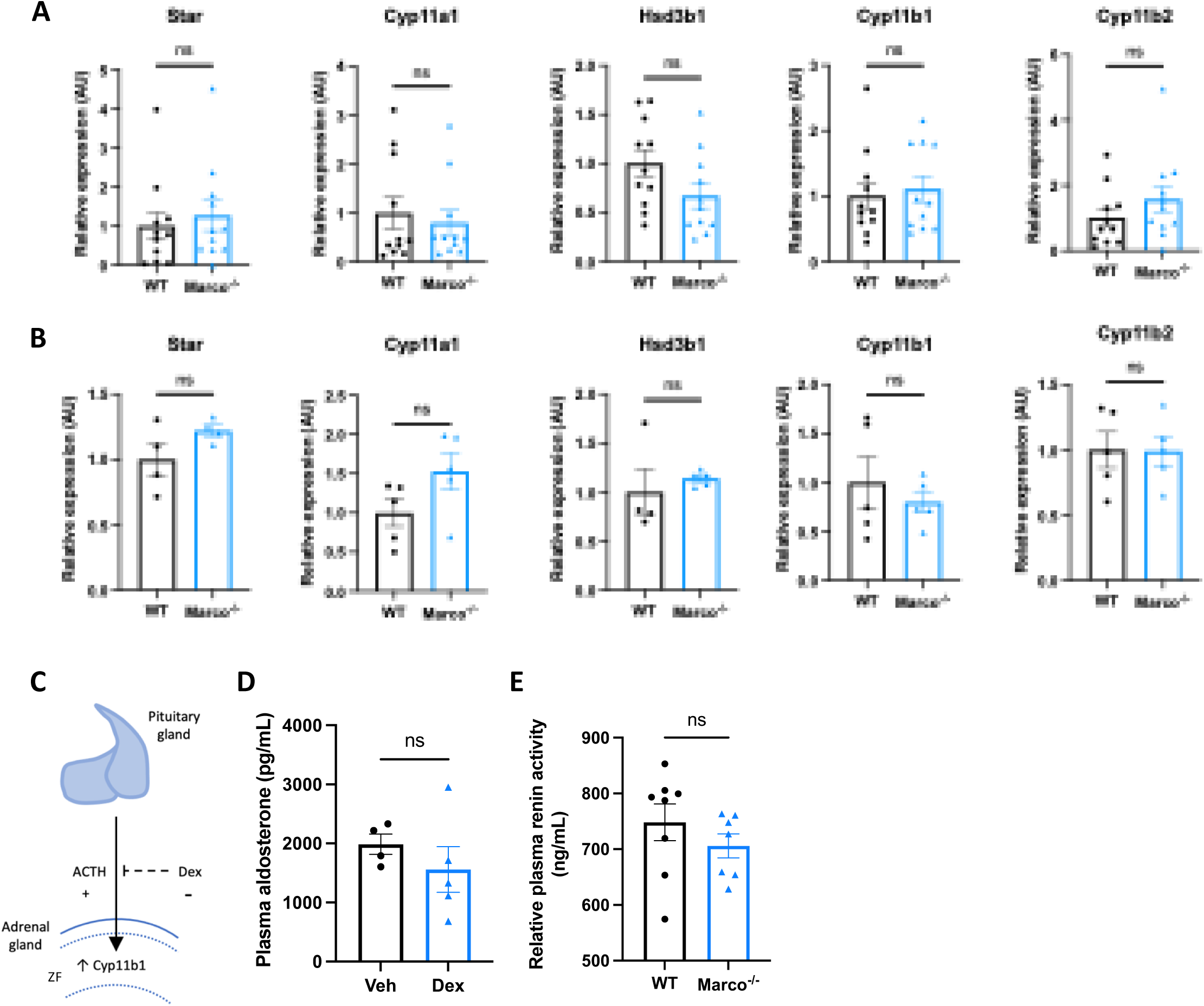
Marco-mediated elevation of aldosterone is not explained by upregulation of adrenal biosynthetic enzymes, the zona fasciculata, or renin. qPCR data reporting the expression of adrenal corticosteroid biosynthetic enzymes between wild type (WT) and Marco-deficient male mice in male (A) and female (B) mice. (C) A schematic illustrating that pituitary-derived adrenocorticotropic hormone (ACTH) stimulates the upregulation of Cyp11b1 from the zona fasciculata (ZF) and the dexamethasone-mediated suppression of this effect. (D) A bar graph showing the plasma aldosterone concentrations of Marco-deficient mice treated with vehicle or dexamethasone-supplemented drinking water for 14 days. (E) A bar graph showing the plasma renin activity of wild type (WT) and Marco-deficient mice at steady state. All data were analysed by two-tailed unpaired Student’s *t*-test and are shown as average ± s.e.m. ns P>0.05. * *P* < 0.05.

CYP11B1 catalyses the conversion of 11-deoxycorticosterone to corticosterone. Corticosterone can be catalysed to aldosterone by CYP11B2. In this sense, the route via CYP11B1 is a bona fide route for the generation of aldosterone, as evidenced by the fact that *Cyp11b1* deletion in mice results in a significant reduction in aldosterone production ^18^. This route is one that is suppressible via the suppressive action of dexamethasone on ACTH and *Cyp11b1* expression. Moreover, ACTH is a known stimulator of aldosterone ^19,20^. Dexamethasone-mediated suppression of the zona fasciculata (Fig. 3c) (Finco et al. 2018) was used to test whether the elevated aldosterone phenotype was zona fasciculata-dependent. Marco-deficient mice fed dexamethasone-supplemented drinking water had plasma aldosterone concentrations comparable to vehicle-treated mice (Fig. 3d), indicating zona fasciculata activity does not contribute to elevated aldosterone levels in Marco^-/-^ mice. Taken collectively, these findings indicate that elevated aldosterone observed in Marco-deficient mice arises extra-adrenally and can therefore be considered a form of secondary hyperaldosteronism. Kidney-derived renin is the initiating hormone in the enzymatic cascade that generates angiotensin II, a potent stimulator of adrenal aldosterone production. We therefore compared plasma renin activity between male WT and Marco^-/-^ mice, finding no significant differences between the two strains (Fig. 3e).

Next, we investigated whether lung-derived angiotensin converting enzyme (*Ace*) could explain the elevated aldosterone levels observed in Marco^-/-^ mice. ACE in the lung catalyses the conversion of Angiotensin I to the aldosterone-stimulating peptide Angiotensin II. We carried out a qPCR test for *Ace* in the lungs of WT and Marco^-/-^ mice in both sexes, finding that Marco-deficient male animals had elevated levels of lung *Ace* relative to WT controls in male mice only (Fig. 4a). Immunofluorescent staining of WT and Marco^-/-^ lungs revealed a substantially higher level of ACE protein in Marco-deficient male mice, while myeloid presence, as measured by CD68 staining, remained unchanged (Fig. 4b). While low levels of ACE expression could be detected in CD68^+^ cells (data not shown), the vast majority of ACE was outside of monocytes and macrophages. We used image analysis software to quantify these changes, finding that ACE median fluorescence intensity (MFI) was significantly increased in male Marco-deficient lungs, while CD68^+^ myeloid cells were present in wild type and knock- out animals at similar levels (Fig. 4c, d). Myeloid cell numbers and lung ACE expression were similar in WT and Marco-deficient female lungs (Fig. 4e, f). We also measured the levels of plasma potassium and sodium levels in male and female WT and Marco-deficient animals, but observed no differences between the genotypes (Fig. 4g-j). Since aldosterone is a known regulator of blood pressure via regulation of blood fluid balance, we also measured blood pressure using the tail-cuff method. We observed that Marco-deficient male mice had marginally reduced diastolic blood pressures, but systolic and mean blood pressure were no different between the two strains (Fig. 4k-m).

**Figure 4.**
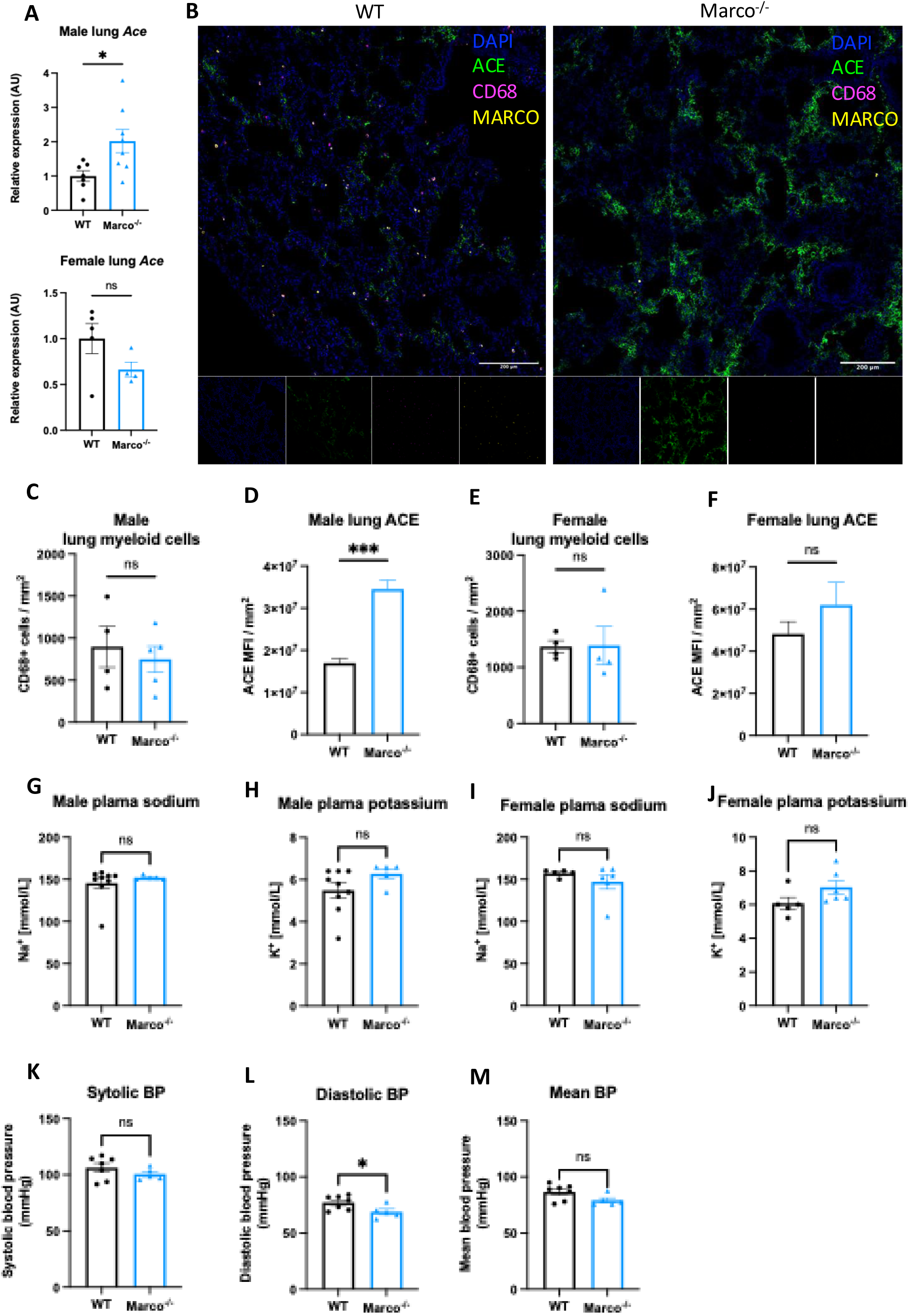
Marco-deficient mice have enhanced expression of Ace in the lung. (A) qPCR data reporting the expression of angiotensin converting enzyme (*Ace*) in the lungs of wild type (WT) and Marco-deficient male and female mice. (B) Representative images showing a cryosection of wild-type and Marco-deficient male murine lungs stained against ACE (green) CD68 (magenta), MARCO (yellow) and DAPI (blue). Quantitation of DAPI-normalised ACE median fluorescence intensity (MFI) (C) and CD68^+^ myeloid cell presence in the lungs of wild type and Marco-deficient male and female mice (C-F). Plasma sodium and potassium levels in male and female mice (G-J). Systolic, diastolic, and mean blood pressure measurements in male mice of the indicated genotypes (K-M). Data in A, C-M, were analysed by two-tailed unpaired Student’s *t*-test and are shown as average ± s.e.m. \**P* < 0.05, \*\**P* < 0.01, \*\*\**P* < 0.001, *\*\*\*\* P* < 0.0001.

We then turned back to analysis of single cell RNA-seq data to identify the cells in the lung that could be mediating this effect. In the lung, unsupervised cluster analysis revealed a total of 12 cell clusters with distinct gene expression signatures (Fig. 5a). We determined the identity of each cell cluster based on the expression of established cell type-specific marker genes, aided by the marker genes identified and outlined in Fig. 5b. We next used dot plots to visualize expression of *Marco* and *Ace* across the different cell clusters. We observed notable *Marco* expression only in alveolar macrophages amongst the different cell clusters (Fig. 5c). *Ace* was shown to be primarily expressed by lung endothelial clusters 1 and 2 (Fig. 5c), in agreement with what is known about lung *Ace* expression. To test whether alveolar macrophages are capable of suppressing endothelial cell *Ace* expression, we co-cultured the MPI alveolar macrophage cell line ^21^, with or without deletion of Marco, with HUVECs endothelial cells for 24 hours, and measured *Ace* expression via qPCR. We observed an increase in *Ace* expression in the Marco-deficient MPI co-cocultures but not WT, though this did not reach statistical significance. Taken collectively, these data suggest a model whereby Marco-expressing alveolar macrophages may, in responses to an as-of-yet unidentified factor, inhibit *Ace* expression at the gene and protein level, and thereby negatively regulate the cleavage of Angiotensin I to form Angiotensin II, and thereby aldosterone production (Fig. 5e).

**Figure 5.**
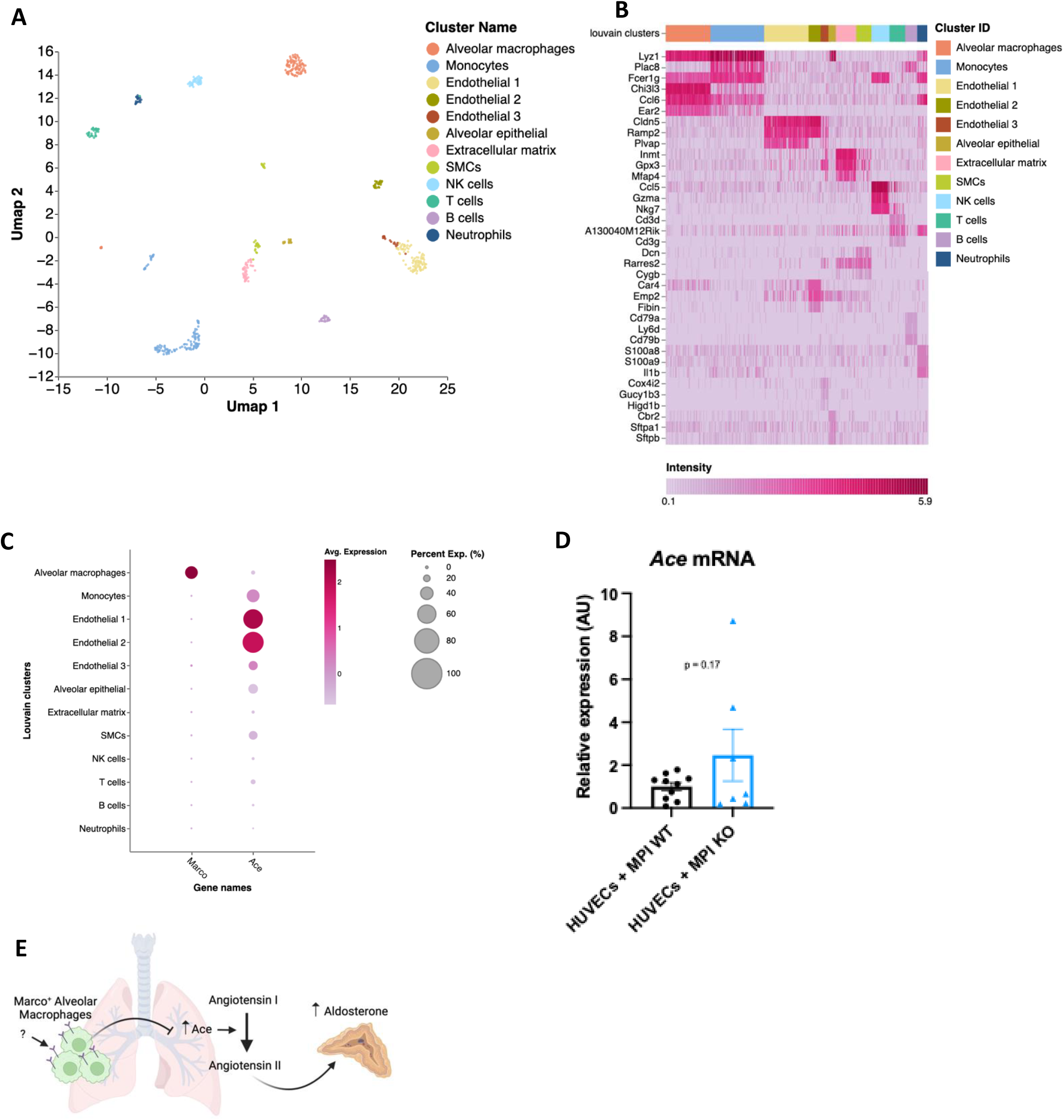
A proposed model for macrophage-mediated regulation of lung Ace expression. (A) Single cell RNA seq UMAP plot depicting the cell types present in the male murine lung. (B) Marker heatmap showing the top three gene markers for each cell cluster in (A). (C) A dot plot showing the gene expression levels of Marco and Ace in the different cell clusters of the male murine lung. (D) qPCR data showing the relative expression levels of Ace in co-coltures contraining HUVECs cells with Marco sufficient or deficient MPI macrophages. (E) A schematic, generated using BioRender, showing the working model by which Marco+ alveolar macrophages regulate aldosterone output from the adrenal gland. Data in D were analysed by two-tailed unpaired Student’s *t*-test and are shown as average ± s.e.m. \**P* < 0.05, \*\**P* < 0.01, \*\*\**P* < 0.001, *\*\*\*\* P* < 0.0001.

In conclusion, we hereby demonstrate that Marco is a negative regulator of aldosterone production, associated with a suppression of angiotensin converting enzyme expression in the lungs of male mice. We propose a model in which extra-adrenal Marco expressing alveolar macrophages, through tissue crosstalk with lung endothelial cells, actively inhibit Ace expression and thereby inhibit the production of aldosterone from the adrenal glands.

## Discussion

Multiple studies over preceding decades have shown that macrophages are much more than simply immune and inflammatory cells. Macrophages have been shown to mediate a wide variety of physiological functions and support homeostasis in virtually every tissue they are found ^22–27^. Our original hypothesis centred on the regulation of corticosteroid output via modulation of cholesterol availability. While we ended up disproving this hypothesis, we did provide a novel example of the non-immune functions of tissue macrophages. It was recently shown that tissue infiltrating macrophages cause neuronal cell death and effective denervation, which disrupts normal melatonin secretion in a model of cardiovascular disease ^28^. To our knowledge, ours is the first study to report that macrophages are involved in the regulation of hormonal output *in vivo* at steady state. It is known that immunosuppressants such as cyclosporine can increase blood pressure ^29^ yet the mechanism remains incompletely understood. This observation is consistent with our data showing macrophage-mediated negative regulation of the blood pressure-controlling hormone aldosterone. Additionally, the fact that higher ACE/ACE2 ratios have been linked to hypertension and worse outcomes in Covid-19 ^30^ could implicate MARCO in the susceptibility to severe disease.

Secondary aldosteronism (SA) is distinguished from primary aldosteronism through renin – SA is characterised by high aldosterone and high or non-suppressed renin levels ^31^. By this definition, Marco-deficient mice exhibit SA, due to their high aldosterone and normal renin levels. SA is also defined as elevated aldosterone that originates outside of the adrenal gland – in this case from ACE in the lung. This finding raises the intriguing possibility that aldosteronism could be triggered or aggravated by MARCO-binding ligands. The fact that blood pressure and plasma electrolytes are not substantially modulated by Marco deletion, despite elevation in plasma aldosterone concentrations, indicate that this mechanism has not evolved to regulate systemic physiology. However, ACE and Angiotensin II are pleiotropic molecules. In addition to their physiological roles, they are also exhibit immunomodulatory functions. *In vitro*, lipopolysaccharide (LPS) treatment of the HUVECs endothelial cell line or human primary endothelial cells causes a reduction in ACE activity, mRNA and protein levels. Similarly, ACE expression in human lung tissue is reduced in sepsis patients relative to control tissues ^32,33^. Supernatant from LPS or TNFα-treated endothelial cells shows a high proportion of ACE^+^ endothelial microparticles (EMPs). ACE^+^ EMPs are also increased in plasma of mice undergoing experimental lung injury and in human septic patients ^33^, indicating shedding of lung endothelial ACE in the context of inflammation. Ace and AngII have been shown to enhance inflammation in a variety of contexts. It may therefore be that Marco-expressing alveolar macrophages exert an immune-modulatory function within the murine lung via controlling tissue *Ace* expression. Alveolar macrophages, which use MARCO to bind and uptake bacteria ^34^, could serve to enhance lung immunity in a pathogen and ACE-dependent manner. Pharmacologic ACE inhibition was previously shown to dampen pro-inflammatory immune cell inflammation in radiation-induced pneumonitis ^35^. Future experiments could therefore explore whether Marco regulates lung tissue Ace expression in the context of inflammation and/or pathogenic challenge. While acute increases of plasma aldosterone have varying effects on endothelial function in humans and animal models, chronic (i.e. genetic) elevations of plasma aldosterone are virtually universally associated with endothelial dysfunction ^36^. This is frequently characterised by reduced endothelial NOS activity from, and acetylcholine action on, endothelial cells ^36^. It would therefore be logical to test whether vascular dysfunction is present in the endothelia of Marco-deficient mice relative to wild type mice.

While the data presented here indicate that Marco deletion is a positive regulator of aldosterone and lung Ace expression, important questions remain to be answered, such as the precise mechanism by which Marco-expressing alveolar macrophages suppress Ace expression, and the ligand(s) responsible for stimulating this effect via the receptor. Additionally, the mechanisms mediating the sex differences observed here remain unexplored. Our findings nevertheless introduce a novel immune paradigm the renin angiotensin aldosterone system.

## Methods

### Mice

C57BL/6J, Marco^-/-^, and hCD68^GFP/GFP^ mice were bred and housed in individually ventilated cages in specific pathogen-free conditions at the University of Oxford. All experiments on mice were conducted according to institutional, national, and European animal regulations. Animal protocols were approved by the animal welfare and ethics review board of the Department of Physiology, Anatomy, and Genetics at the University of Oxford.

### Tissue collection

All tissues were collected from 8-11 week old male mice that were culled between 0930-1100. Mice were euthanised by intraperitoneal injection of 10μL/g 200mg/mL Pentoject (pentobarbital sodium; Animalcare). Following cessation of the pedal motor reflex a sample of blood was taken from the right atrium prior to perfusion through the left ventricle with 20mL 1X phosphate-buffered saline (PBS; Sigma Aldrich). Adrenal glands were dissected, and the peri-adrenal fat removed using a stereomicroscope. Following harvest, adrenal glands and lungs were frozen at -80°C until subsequent molecular analyses or fixed overnight in 4% paraformaldehyde (PFA) prior to histological analyses. Bloods obtained prior to perfusion were collected into EDTA coated tubes to prevent clotting and centrifuged at 1000g for 15 minutes to separate plasma. Plasma samples were frozen at -80°C in 1.5mL Eppendorf tubes until subsequent analyses.

### Plasma hormone, cholesterol, renin activity, and electrolyte quantitation

We measured plasma aldosterone and corticosterone using reverse ELISA kits (Enzo Life Sciences) according to manufacturer’s instructions. Plasma cholesterol was quantified by colorimetric assay (Cholesterol Quantitation Kit; Sigma Aldrich) according to manufacturer’s instructions. Plasma renin activity was measured by fluorometric assay using the Renin Assay Kit (Sigma Aldrich). Colorimetric and fluorimetric readouts from these assays were attained using a FLUOstar Omega plate reader (BMG Labtech). Plasma electrolytes were outsourced to Pinmoore Animal Laboratory Services Limited (Cheshire, England) and sodium and potassium electrolyte readings were taken using an I-Smart 30 Vet electrolyte analyser (Woodley Equipment).

### Single cell sequencing data analysis

Publicly available single cell RNA-seq datasets (steady-state adrenal: PMID 33571131, GEO accession number GSE161751; steady-state lung: PMID 30283141, GEO accession number GSE109774) were processed, explored and visualised using Cellenics® community instance (https://scp.biomage.net/) hosted by Biomage (https://biomage.net/). For the adrenal dataset the classifier filter was disabled as the sample had been pre-filtered. The cell size distribution filter was disabled. A mitochondrial content filter was used, with the absolute threshold method used (bin step = 0.05, maximum fraction = 0.1). The number of genes vs UMI filter was used with a linear fit and P value = 0.0004906771. The doublet filter was used (bin step = 0.05, probability threshold = 0. 8743427. The harmony algorithm was used for data integration (2000 genes, log normalisation). The RPCA method was used for dimensionality reduction (13 principal components). Cells and clusters were visualised using UMAP embedding with the cosine distance metric (minimum distance = 0.2). Louvain clustering was the clustering algorithm used in this analysis (resolution = 0.6). For the lung dataset the classifier filter was disabled as the sample had been pre-filtered. A cell size distribution filter was utilised with binStep = 200 and minimum cell size = 1669. A mitochondrial content filter was used, with the absolute threshold method used, bin step = 0.3, maximum fraction = 0. The number of genes vs UMI filter was used with a linear fit and P value = 0.001. The doublet filter was used with bin step = 0.02 and probability threshold = 0.7616616. The harmony algorithm was used for data integration (2000 genes, log normalisation). The RPCA method was used for dimensionality reduction (24 principal components). Cells and clusters were visualised using UMAP embedding with the cosine distance metric (minimum distance = 0.3). Louvain clustering was the clustering algorithm used in this analysis (resolution = 0.8).

### Cryo-sectioning and antibody staining

Prior to tissue embedding, adrenal glands were cryoprotected in 30% sucrose solution (in PBS) overnight. Cryoprotected adrenal glands were embedded in OCT embedding medium (Thermo Fisher Scientific) and snap frozen in liquid nitrogen. 15um sections were cut from embedded tissues using a Leica cryostat and mounted onto charged microscope slides. Cryosectioned tissue slices were thawed in PBS for 10 min at room temperature, tissue sections were circled using a super PAP pen (Life Technologies), blocked and permeabilised for 1 hour at room temperature in perm/block solution (3% bovine serum albumin, 2% goat serum, 1% Triton X-100, 0.01% NaN3 in PBS). Slides were stained with CD68-AF647 (clone FA-11, BioLegend), chicken anti-GFP (ab13970, Abcam) with goat anti-chicken AF488 (A11039, Invitrogen), mouse anti-MARCO (ED31, BioRad) with goat anti-mouse AF546 (A-11030, Invitrogen), and rabbit anti-ACE (MA5-32741, Invitrogen) with goat anti-rabbit Alexa Fluor 488 Tyramide Super Boost kit (Invitrogen, B40922). Primary stains were done overnight at 4°C and secondary stains were done for 1 hour at room temperature. The goat anti-rabbit Alexa Fluor 488 Tyramide Super Boost kit was used as per manufacturer’s instructions. DAPI counterstains were done at 1:1000 for 5 minutes at room temperature. Fluoromount G mounting medium (Invitrogen) was used to mount coverslips to slides which were then sealed with clear nail varnish. Immunofluorescence images were acquired using the Zeiss 880 confocal microscope and images were analysed in FIJI.

### RT-PCR

Tissues were homogenised using the Precellys Hard Tissue Grinding Kit (MK28-R; Bertin Technologies). Total RNA from homogenised adrenals or lungs was isolated using RNeasy Plus Micro Kit (Qiagen, cat# 74034). cDNA was reverse transcribed using SuperScript II (Invitrogen) and random primers (Invitrogen). Quantitative PCR was performed using SYBR Green (Applied Biosystems) in C1000 Touch™ Thermal Cycler (BioRad). Β-actin was used as the housekeeping gene to normalize samples. We used the following formula to calculate the relative expression levels: RQ = 2^^−ΔCt^ × 100 = 2^−(Ct gene of interest – Ct β actin)^ × 100. The following mouse-specific primers were used. *Star* forward 5’-CAGGGCCAAGAAAACCTACA-3’; *Star* reverse 5’-ACGAGCATTTTGAAGCACCT-3’; *Cyp11a1* forward 5’-AGGACTTTCCCTGCGCT -3’; *Cyp11a1* reverse 5’-GCATCTCGGTAATGTTGG-3’; *Hsd3b1* forward 5’-GCGGCTGCTGCACAGGAATAAAG-3’; *Hsd3b1* reverse 5’-TCACCAGGCAGCTCCATCCA-3’; *Cyp11b1* forward 5’-TCACCATGTGCTGAAATCCTTCCA-3’; *Cyp11b1* reverse 5’-GGAAGAGAAGAGAGGGCAATGTGT-3’; *Cyp11b2* forward 5’-CAGGGCCAAGAAAACCTACA-3’; *Cyp11b2* reverse 5’-ACGAGCATTTTGAAGCACCT-3’; *Β-actin* forward 5’-TCATGAAGTGTACGTGGACATCC-3’; *Ace* forward 5’-GCTTCCTCTTTCTGCTGCTCTG -3’; *Ace* reverse 5’- TGCCCTCTATGGTAATGTTGGT- 3’; B*-actin* reverse 5’-CCTAGAAGCATTTGCGGTGGACGATG-3’.

### Zona fasciculata suppression model

Water-soluble dexamethasone (Sigma-Aldrich) was reconstituted in autoclaved deionized water at the concentration of 0.0167 mg/mL as described in ^37^. 10 week old Marco^-/-^ mice were fed dexamethasone-supplemented drinking water for 14 days prior to blood sampling as described above.

### Digital image analysis

Quantitation of the area and intensities of immunofluorescent stains was done using FIJI image analysis software. Three cryosections of stained lung tissue (∼5mm^2^ tissue acquired per section) were analysed per mouse. CD68^+^ cells were segmented using the Otsu thresholding plugin. The parameters were adjusted manually to ensure optimum cell segmentation. DAPI-stained nuclei were segmented using the Otsu thresholding plugin. Median fluorescence intensity (MFI) for ACE was measured via the integrated density of the stain, which was normalised to the area of DAPI staining.

### Blood pressure readings

Systolic, diastolic, and mean blood pressures were taken using the CODA high throughput noninvasive blood pressure system (Kent Scientific). Mice were acclimatised to animal holders on three occasions in the week prior to experimental readings.

### Co-culture

X63-GMCSF and WT and Marco-deficient MPI cells were a kind gift from Dr Subhankar Mukhopadhyay. 20,000 HUVECs cells were plated with 20,000 WT or Marco-deficient MPI cells in 96 well plates for 24 hours in high glucose DMEM, 10% Fetal Bovine Serum (FBS), 1% X63-GM-CSF conditioned media as a source of GM-CSF, 1% Penicillin/Streptomycin (P/S), and 1X MesoEndo Cell growth medium (all bought from Sigma Aldrich) and incubated at 37°C, 5% CO_2_. RNA was extracted and qPCRs performed as previously described.

### Statistical Analysis

Results are expressed as the mean ± s.e.m., as indicated in the figure legends. Statistical significance between two experimental groups was assessed using two-tailed student’s t test. All Statistical analyses were performed in GraphPad Prism 9 (GraphPad, USA) for Mac OS X. All calculated P values are reported in the figures, denoted by \**P* < 0.05, \*\**P* < 0.01, \*\*\**P* < 0.001, ns P>0.05.

## References

1. Shi, B. et al. The Scavenger Receptor MARCO Expressed by Tumor-Associated Macrophages Are Highly Associated With Poor Pancreatic Cancer Prognosis. Front Oncol 11, 4518 (2021).

2. Van Der Laan, L. J. W. et al. Macrophage scavenger receptor MARCO: In vitro and in vivo regulation and involvement in the anti-bacterial host defense. Immunol Lett 57, 203–208 (1997).

3. Arredouani, M. et al. The Scavenger Receptor MARCO Is Required for Lung Defense against Pneumococcal Pneumonia and Inhaled Particles. Journal of Experimental Medicine 200, 267–272 (2004).

4. Xing, Q. et al. Scavenger receptor MARCO contributes to macrophage phagocytosis and clearance of tumor cells. Exp Cell Res 408, 112862 (2021).

5. Kanno, S. et al. Scavenger receptor MARCO contributes to cellular internalization of exosomes by dynamin-dependent endocytosis and macropinocytosis. Scientific Reports 2020 10:*1* 10, 1–12 (2020).

6. Eisinger, S. et al. Targeting a scavenger receptor on tumor-associated macrophages activates tumor cell killing by natural killer cells. doi:10.1073/pnas.2015343117/-/DCSupplemental.

7. Plüddemann, A., Neyen, C. & Gordon, S. Macrophage scavenger receptors and host-derived ligands. Methods 43, 207–217 (2007).

8. Rigotti, A. et al. A targeted mutation in the murine gene encoding the high density lipoprotein (HDL) receptor scavenger receptor class B type I reveals its key role in HDL metabolism. Proc Natl Acad Sci U S A 94, 12610 (1997).

9. Temel, R. E. et al. Scavenger receptor class B, type I (SR-BI) is the major route for the delivery of high density lipoprotein cholesterol to the steroidogenic pathway in cultured mouse adrenocortical cells. Proc Natl Acad Sci U S A 94, 13600–13605 (1997).

10. Hoekstra, M. et al. Scavenger receptor class B type I-mediated uptake of serum cholesterol is essential for optimal adrenal glucocorticoid production. J Lipid Res 50, 1039 (2009).

11. Ito, M. et al. SR-BI (Scavenger Receptor BI), not LDL (Low-Density Lipoprotein) receptor, mediates adrenal stress response - Brief report. Arterioscler Thromb Vasc Biol 40, 1830–1837 (2020).

12. Santos, R. A. S. et al. The renin-angiotensin system: Going beyond the classical paradigms. Am J Physiol Heart Circ Physiol 316, H958–H970 (2019).

13. Baudrand, R. et al. Statin Use and Adrenal Aldosterone Production in Hypertensive and Diabetic Subjects. Circulation 132, 1825–1833 (2015).

14. Hornik, E. S. et al. A clinical trial to evaluate the effect of statin use on lowering aldosterone levels. BMC Endocr Disord 20, (2020).

15. Simpson, H. D. et al. Effects of cholesterol and lipoproteins on aldosterone secretion by bovine zona glomerulosa cells. Journal of Endocrinology 121, 125–131 (1989).

16. Cherradi, N. et al. Angiotensin II Promotes Selective Uptake of High Density Lipoprotein Cholesterol Esters in Bovine Adrenal Glomerulosa and Human Adrenocortical Carcinoma Cells Through Induction of Scavenger Receptor Class B Type I. Endocrinology 142, 4540–4549 (2001).

17. Kopprasch, S. et al. Prediabetic and diabetic in vivo modification of circulating low-density lipoprotein attenuates its stimulatory effect on adrenal aldosterone and cortisol secretion. Journal of Endocrinology 200, 45–52 (2009).

18. Mullins, L. J. et al. Cyp11b1 null mouse, a model of congenital adrenal hyperplasia. J Biol Chem 284, 3925–3934 (2009).

19. Seely, E. W., Conlin, P. R., Brent, G. A. & Dluhy, R. G. Adrenocorticotropin stimulation of aldosterone: prolonged continuous versus pulsatile infusion. J Clin Endocrinol Metab 69, 1028–1032 (1989).

20. Daidoh, H. et al. Responses of plasma adrenocortical steroids to low dose ACTH in normal subjects. Clin Endocrinol (Oxf) 43, 311–315 (1995).

21. Fejer, G. et al. Nontransformed, GM-CSF-dependent macrophage lines are a unique model to study tissue macrophage functions. Proc Natl Acad Sci U S A 110, E2191 (2013).

22. Theurl, I. et al. On-demand erythrocyte disposal and iron recycling requires transient macrophages in the liver. Nature Medicine 2016 22:8 22, 945–951 (2016).

23. Cox, N. et al. Diet-regulated production of PDGFcc by macrophages controls energy storage. Science (1979) 373, eabe9383 (2021).

24. Muller, P. A. et al. Crosstalk between Muscularis Macrophages and Enteric Neurons Regulates Gastrointestinal Motility. Cell 158, 300–313 (2014).

25. Hulsmans, M. et al. Macrophages Facilitate Electrical Conduction in the Heart. Cell 169, 510–522.e20 (2017).

26. Schafer, D. et al. Microglia sculpt postnatal neural circuits in an activity and complement-dependent manner. Neuron 74, 691–705 (2012).

27. Pirzgalska, R. M. et al. Sympathetic neuron-associated macrophages contribute to obesity by importing and metabolizing norepinephrine. Nat Med 23, 1309–1318 (2017).

28. Ziegler, K. A. et al. Immune-mediated denervation of the pineal gland underlies sleep disturbance in cardiac disease. Science (1979) 381, 285–290 (2023).

29. Robert, N., Wong, G. W. & Wright, J. M. Effect of cyclosporine on blood pressure. Cochrane Database of Systematic Reviews (2010) doi:10.1002/14651858.CD007893.PUB2/MEDIA/CDSR/CD007893/IMAGE_N/NCD007893-CMP-003-09.PNG.

30. Pagliaro, P. & Penna, C. ACE/ACE2 Ratio: A Key Also in 2019 Coronavirus Disease (Covid-19)? Front Med (Lausanne) 7, 335 (2020).

31. Pócsai, K., Sumánszki, C. & Tőke, J. Secondary Aldosteronism. Practical Clinical Endocrinology 309–317 (2021) doi:10.1007/978-3-030-62011-0_29.

32. Hermanns, M. I., Müller, A. M., Tsokos, M. & Kirkpatrick, C. J. LPS-induced effects on angiotensin I-converting enzyme expression and shedding in human pulmonary microvascular endothelial cells. In Vitro Cell Dev Biol Anim 50, 287–295 (2014).

33. Takei, Y. et al. Increase in circulating ACE-positive endothelial microparticles during acute lung injury. European Respiratory Journal 54, 1801188 (2019).

34. Palecanda, A. et al. Role of the Scavenger Receptor MARCO in Alveolar Macrophage Binding of Unopsonized Environmental Particles. Journal of Experimental Medicine 189, 1497–1506 (1999).

35. Sharma, G. P. et al. Pharmacologic ACE-Inhibition Mitigates Radiation-Induced Pneumonitis by Suppressing ACE-Expressing Lung Myeloid Cells. International Journal of Radiation Oncology*Biology*Physics 113, 177–191 (2022).

36. Toda, N., Nakanishi, S. & Tanabe, S. Aldosterone affects blood flow and vascular tone regulated by endothelium-derived NO: therapeutic implications. Br J Pharmacol 168, 519 (2013).

37. Finco, I., Lerario, A. M. & Hammer, G. D. Sonic Hedgehog and WNT Signaling Promote Adrenal Gland Regeneration in Male Mice. Endocrinology 159, 579 (2018).

